# Elevated [CO_2_] concentration and nitrogen addition affects responses of foliar phosphorus fractions in invasive species to increased phosphorus supply

**DOI:** 10.1101/2021.01.03.425118

**Authors:** Lingling Zhang, Xianzhen Luo, Hans Lambers, Guihua Zhang, Nan Liu, Xiaowei Zang, Meijuan Xiao, Dazhi Wen

**Affiliations:** CAS Key Laboratory of Vegetation Restoration and Management of Degraded Ecosystems, South China Botanical Garden, Chinese Academy of Sciences, Guangzhou 510650, China; Southern Marine Science and Engineering Guangdong Laboratory, Guangzhou 511458, China; School of Biological Sciences, The University of Western Australia, Crawley (Perth), WA 6009, Australia; Department of Plant Nutrition, College of Resources and Environmental Sciences, National Academy of Agriculture Green Development, Key Laboratory of Plant–Soil Interactions, Ministry of Education, China Agricultural University, Beijing 100193, China; CAS Engineering Laboratory for Vegetation Ecosystem Restoration on Islands and Coastal Zones, South China Botanical Garden, Chinese Academy of Sciences, Guangzhou, 510650, China

**Keywords:** Phosphorus availability gradient, Elevated CO_2_, Photosynthetic rates, Foliar P fractions, Invasive plant species

## Abstract

No studies have explored how the invasive species of *Mikania micranatha* and *Chromolaena odoratan* adjust leaf phosphorus (P) among inorganic P (Pi) and organic P fractions to adapt the low soil P availability, especially under elevated CO_2_ concentrations ([CO_2_]) and nitrogen (N) deposition. Here, we address this by measuring foliar total N and P concentrations as well as functional P fractions (i.e. Pi, metabolic P, lipid P, nucleic acids P, and residual P) of both invasive species and a native species (*Paederia. scandens*) growing under different P supplies, N, and N+P addition under both ambient and elevated [CO_2_]. Phosphorus addition greatly increased plant biomass and foliar P concentrations but did not significantly affect foliar N concentration and leaf mass per unit leaf area (LMA). In response to P addition, the concentration of metabolic P increased the most, followed by that of nucleic acid P, Pi, and lipid P, in all species by an average of 754%, 82%, 53%, and 38%, respectively. However, elevated [CO_2_] and N addition weakened this positive effect on concentrations of foliar P fractions in the invasive species. Our results indicate that elevated [CO_2_] and N addition allowed the invasive species to acclimate to a low soil P availability, supporting their successful invasion, through greatly reducing P allocation to non-metabolic foliar P fractions (phospholipids and nucleic acid P) to meet their demand for metabolic P and Pi for photosynthesis, rather than altering LMA.

## Introduction

Phosphorus (P) is an essential plant nutrient and often present in soil at concentrations that limit net primary productivity (Hidaka and Kitayama, 2013; Mo et al., 2019). In subtropical forests, plant productivity is generally limited by a low availability of soil P, rather than by a low availability of nitrogen (N) due to the long-term weathering of bedrock and the gradual loss of P (Walker and Syers, 1976; Jonard et al., 2015; Mo et al., 2019). While P is often limiting, N is increasingly available in subtropical forests because of atmospheric N deposition, which has increased to ~ 30-50 kg N ha year^−1^ in subtropical forests in China (Mo et al., 2006; Luo et al., 2019). Another factor that may greatly affect plant growth is the atmospheric concentration of CO_2_ ([CO_2_]), which has increased from ~280 μmol mol^−1^ in 1840s to ~ 410 μmol mol^−1^ in 2020 (IPCC, 2013; Luo et al., 2019; https://www.co2.earth/). However, the effects of increases in N deposition and atmospheric [CO_2_] on the strategies that plants have evolved to use P efficiently in P-impoverished forests are rarely documented in invasive species (Campbell and Sage, 2006; Lewis et al., 2010; Tissue and Lewis, 2010). Invasive species threaten plant diversity (Dukes and Mooney, 1999), and can potentially alter the function and structure of terrestrial ecosystems (Li et al., 2002; Tang et al., 2007; Song et al., 2009; Sage, 2019). Understanding how the strategies of allocation of foliar P for maintaining plant productivity in invasive species is affected by N deposition and elevated [CO_2_] would increase the ability to predict and perhaps control plant invasions in tropical P limited forest ecosystems (Song et al., 2009; Wang et al., 2016).

Regulating P allocation to leaves is a vital strategy in plants to acclimate to soil conditions (e.g., low soil P availability) (Zhang et al., 2018; Wang et al., 2019) and climate change (e.g., elevated [CO_2_] and N deposition) (Tissue and Lewis, 2010). Foliar P can be fractionated into inorganic phosphate (Pi) and organic P fractions (metabolic P, lipid P, nucleic acid P and residuals P) (Hidaka and Kitayama, 2013). Pi represents a significant fraction of leaf P, and is generally stored in the vacuole when a plant is unable to acquire adequate Pi from the soil (Veneklaas et al., 2012). The foliar metabolic P fraction consists mainly of intermediates of carbon metabolism, such as bioactive-molecular compound (e.g., phosphorylated sugars, ADP and ATP). The major organic P fraction, i.e. nucleic acid P, often represents more than 50% of the foliar organic P pool; up to 85 % of nucleic acid P consists of rRNA, which is essential for protein synthesis (Matzek & Vitousek, 2009). Lipid P comprises phospholipids, most of which are components of the plasmalemma and organelle membranes (Veneklaas et al., 2012). Finally, the uncharacterized residual fraction may include phosphorylated proteins, some of which regulate cellular processes (Yan et al., 2019). Despite studies on allocation of leaf P fractions following N or P addition (Mo et al., 2019) and in different soil condition (e.g., soil age; Yan et al., 2019), there is little information concerning the interactive effects of elevated [CO_2_], N addition, and low soil P availability on the allocation of P to the five foliar P fractions in invasive species (Song et al., 2009; Tissue and Lewis, 2010; Zhang et al., 2016).

Under P deficiency, photosynthesis is generally reduced, due to feedback inhibition resulting for reduced leaf growth (Zhang et al., 2016) or the limitation of orthophosphate (Pi) in the cytosol (Mo et al., 2019). These decreases in photosynthetic activity might increase photosynthetic N-use efficiency (PNUE) and photosynthetic P-use efficiency (PPUE), and also decrease the leaf mass per unit leaf area (LMA; Ghannoum et al., 1999). Plant grow in low soil P availability can reduce their overall need for foliar P by decreasing metabolic P fractions, and buffer direct Pi restriction of photosynthesis (Hadiaka & Kitayama, 2011; Warren, 2011). Moreover, the replacement of phospholipids (lipid P) in membranes by sulfolipids and galactolipids to maintain foliar metabolic P concentration in P-deficiency soil (Lambers et al., 2012; Veneklass et al., 2012). For invasive plants growing in soils with low P availability, however, how they adapt the low soil P availability under elevated [CO_2_] and N addition remain unclear.

Since their invasion of southern China in the 1980s, *Mikania micranatha* and *Chromolaena odorata* have caused serious damage to secondary forests and crops (Li and Xie, 2002; Song et al., 2009). The rapid spread of both invasive plants has triggered a serious decline in the diversity of native species in terrestrial ecosystems (D’Antonio et al., 2004; Bradley et al., 2010). The photosynthetic rate is faster in invasive species than in co-occurring native species (Baruch and Goldstein, 1999; Deng et al., 2004: Song et al., 2009). Relative to native species, invasive species generally have greater phenotypic plasticity, are more tolerant to environmental change, such as elevated [CO_2_], N deposition, or low soil P availability (Alpert et al., 2000; Geng et al., 2006; Feng et al., 2007; Tissue and Lewis, 2010).

The objectives of this study were: 1) to determine how the invasive plants *M. micrantha* and *C. odorata* respond to low P availability (in terms of P allocation to leaves and related foliar traits) in order to maintain photosynthetic rates and 2) to determine how those responses are affected by elevated [CO_2_], and N deposition. To accomplish these objectives, we conducted an open-top field chamber experiment. We hypothesized that (1) foliar traits (i.e. LMA and N and P concentrations) and the photosynthetic capacity of the invasive species would increase with increasing P-application rate, and that these increases would be greater with elevated [CO_2_] than with N addition; (2) the increase in photosynthetic capacity in response to P and N addition under elevated [CO_2_] would be more pronounced in invasive species than in a native species; and (3) elevated [CO_2_] and N addition would change the pattern of allocation of P to foliar P fractions for photosynthesis and thereby allow the invasive plants to maintain plant growth in a soil with low P availability.

## Materials and methods

### Site description

The open-top field chamber experiment was conducted at South China Botanical Garden (23°08′N, 113°17′E), located in Guangzhou Province, China. The region has a subtropical monsoon climate (Zhang et al., 2016; Luo et al., 2019) with a mean annual precipitation of 1750 mm, a mean annual temperature of 21.5 °C, and a mean relative air humidity of 77% (Zhang et al., 2016).

### Experimental design

The experiment included two widespread invasive species, i.e., *M. micranatha* and *C. odorata*. For comparison, the experiment also included a native species that has a similar morphologies as the invasive species, i.e. *Paederia scandens*. *M. micrantha*, *C. odorata* and *P. scandens* were collected in South China Botanical Garden.

Seedlings were initially grown under suitable soil water and light conditions in a nursery. Seedlings of similar size (about 100 mm tall) were then transplanted into pots (one seedling per pot) that were 280-mm tall and 320-mm in diameter and contained 20 kg of soil. The soil had been collected at 0-400 mm depth from a primary broadleaf forest in South China Botanical Garden; the soil was mixed before it was transferred to the pots. The soil chemical properties (means ± SE) before treatments were: pH= 5.0±0.05; organic C = 16.1±0.6 mg g^−1^; total N = 1.9±0.04 mg g^−1^; total P = 0.35±0.02 mg g^−1^; NH_4_-N = 30±3.1 mg kg^−1^; and NO_3_-N = 8.1±0.2 mg kg^−1^. Each species was represented by 120 pots.

The experiment used 12 open-top chambers. Six of the chambers were “new”, i.e., they were constructed in January 2016, and were cylindrical, 3.5 m in height, and 5.0 m in diameter. The other six open-top chambers were “old”, i.e., they were constructed in 2012, and were cylindrical, 3.5 m in height, and 3.4 m in diameter. Details on the construction of chambers and on the method of supplying CO_2_ were described previously (Zhang et al., 2016).

On 18 June 2016, uniform and healthy seedlings of each species were selected and assigned to each chamber; each chamber contained 18 pots (for the narrow chambers) or 24 pots (for the wide chambers) of each species (Fig. S1). On 2 July 2016, two old and four new open-top chambers were exposed to elevated [CO_2_] (700±50 μmol mol^−1^), and the other six chambers were exposed to ambient [CO_2_] (400±50 μmol mol^−1^) (Fig. S1). We detected no significant difference in the light, temperature, or moisture conditions between the old and new chambers. For each species in each wider chamber, four pots were not treated with P or N and were used as controls; four pots were treated with N (6.25 g N m^−2^ yr^−1^); four pots each were treated with P fertilizer at rates of 0.75 (1/2P), 1.5 (1P), or 3.0 (2P) g P m^−2^ yr^−1^; and four pots were treated with both N and P (6.25 g of N m^−2^ yr^−1^ + 1.5 g of P m^−2^yr^−1^). In total, there were 12 conditions: three levels of P addition, one level of N addition, one level of N+P addition, the control, and two levels of CO_2_ for each of the previous six conditions. For each species in each new chamber, the same treatments were applied to three rather than to four pots per species. Chemically pure NH_4_NO_3_ was used as the N source, and chemically pure NaH_2_PO_4_·2H_2_O was used as the P source (Guangzhou Chemical Reagent Factory, Guangzhou, China). The solutions of N, P, or NP were sprayed on the soil surface of each pot.

### Measurement of foliar gas exchange

Healthy sun-exposed mature leaves were chosen for foliar gas exchange measurement from 9: 00 am to 12: 00 am during 11 days in October 2016. For each gas exchange coefficient, at least six individuals for each combination of species and treatment were measured. Following the order of photosynthetic photon flux density (PPFD) 1200, 1000, 800, 500, 300, 200, 120, 50, 20, 0 μmol m^−2^ s^−1^, photosynthetic light-response curves were made. When the measurements were conducted, the vapor pressure deficit was set at 2.0±0.5 kPa, and leaf temperature at 30±1 °C. The nonrectangular hyperbola model of Thornley (1976) was used to calculate the maximum light-saturated photosynthetic rate.

### Measurement of foliar structural traits

After leaf gas exchange was measured, the leaf area of each projected leaf was determined using a leaf area meter (LI-3100C; LI-COR Biosciences, Nebraska, USA); these leaves were then collected and oven-dried at 65 ◻ to a constant weight for calculating the leaf mass per area (LMA). The remaining mature leaves on the sampled branch of each plant were freeze-dried and ground (after main veins and petioles were removed) to determine their foliar N and P concentrations, and the concentrations of foliar P fractions. Foliar N and P concentrations were measured by a colorimetric assay after sulfuric acid (H_2_SO_4_) digestion (Sommers et al., 1970; Carter and Gregorich, 2007; Liu et al., 1996).

PPUE and PNUE were calculated as the ratio of the maximum photosynthetic rates per unit P or N. Photosynthetic capacity was expressed on a leaf area and dry mass basis. Finally, the plants were harvested, and divided into roots, stems and leaves, then dried and weighed to calculate the biomass.

### Measurement of leaf P fractions

Foliar P is generally divided into inorganic P (Pi) and organic P (metabolic P, lipid P, nucleic acid P, and residual P). Organic P fractions were sequentially extracted (Hidaka and Kitayama, 2013) following methods of Kedrowski (1983) and Close & Beadle (2004). Foliar Pi was extracted by the acetic-acid extraction method (Yan et al., 2019), and determined using a molybdenum blue-based method (Ames, 1966). The four fractions of organic P were determined in the following steps. First, a 0.5-g subsample of a freeze-dried and ground foliar sample was homogenized with 15 ml of 12: 6: 1 CMF (chloroform, methanol, and formic acid, v/v/v) in a 50-ml centrifuge tube (first tube). The liquid was extracted twice with a total of 19 ml of 1: 2: 0.8 CMW (chloroform, methanol, water, v/v/v), and added with 9.5 ml of chloroform-washed water. The final solvent was 1:1: 0.9 CMW (v/v/v), which caused the extract to separate into a sugar-and nutrient-rich upper layer and a lipid-rich organic bottom layer. The upper layer in the second tube was transferred to a new tube (the third tube), and the bottom layer was used to determine lipid P.

A 5-ml volume of 85% methanol (v/v) was added to the material in the third tube, which was then placed in a vacuum dryer for 48 h to remove dissolved chloroform and methanol. The aqueous layer was refrigerated (4 °C) for 1 hr, and 5 % trichloroacetic acid (TCA) solution was made through adding 1 ml 100 % (w/v) TCA. A 10-ml volume of cold 5 % (w/v) TCA was then added to the tube. After 1 hr, the material in the tube was shaken for 1 hr and then centrifuged at 3000 g for 10 min. The supernatant was prepared for the determination of the sum of Pi and metabolic P. We subtracted Pi from the sum to obtain the metabolic P.

Finally, the remaining residue after extraction of the cold TCA was mixed with 35 ml 2.5 % TCA (w/v), and extracted for 1 hr at 95 °C in a hot water bath. Aliquots were centrifuged at 3000 g for 10 min, and taken for analysis of nucleic acid P. The residue remaining from the hot TCA final extraction was the residual P fraction. The determination method of all foliar P fractions was similar to that of foliar total P, and the quantity of the fractions were expressed on a dry mass basis.

### Data analyses

The foliar N to total P ratio or the foliar N to P fraction ratios were calculated based on mass. The effects of species, N addition, P addition, elevated [CO_2_] and their interactions on foliar P fractions and foliar traits were assessed by multi-way ANOVA with species, N and P addition, and elevated [CO_2_] as fixed factors. One-way analyses of variance (ANOVAs) were used to compare the effects of treatments on LMA, PPUE, PNUE, N: P ratios, and the concentrations of N, P, and P fractions. Relationship between plant biomass and concentration of foliar P fractions, total P was conducted with Regression linear analysis. Means were compared with LSD test at significance level of 0.05. SPSS 19.0 (SPSS. Inc., Chicago, IL., USA) was used for statistical analyses, and figures were drawn with Origin 2015 (Origin Lab. Inc., Massachusetts, USA).

## Results

### Photosynthetic capacity (A_area_ and A_mass_) and leaf mass per area (LMA), and plant biomass

Elevated [CO_2_], N addition, species, and their interactions significantly affected photosynthetic capacity (A_area_), but P addition did not significantly affect photosynthetic capacity (A_area_ and A_mass_) in invasive and native species (Table 1). Elevated [CO_2_], N addition, species, and P addition significantly affected plant biomass, but their interactions did not significantly affect plant biomass (Table 1). 2P addition significantly increased plant biomass of *M. micrantha*, *C. odorata*, *and P. scandens* by 8, 43, and 28%, respectively (Fig. 1). Elevated [CO_2_] significantly increased A_area_ and A_mass_ in the invasive species (*M. micrantha* and *C. odorata*) by 17-101% and 5.5-97%, respectively (Fig. 2d, e, g, h); however, elevated [CO_2_] did not significantly affect A_area_ or A_mass_ in the native species (*P. scandens*) (Fig. 2f, i). N addition also significantly increased A_area_ and A_mass_ in *M. micrantha* and *C. odorata*, by 35-38% and 2.8-41%, respectively, under elevated [CO_2_] (Fig. 2d, e, g, h), but did not significantly affect A_area_ or A_mass_ in *P. scandens*. Elevated [CO_2_], N addition, and P addition did not significantly affect LMA in *M. micrantha* or *C. odorata*, while 2P addition increased LMA significantly more than 1P addition in *P. scandens* (Table 1 and Fig. 2a, b, c).

**Table 1.** Effects of species (S), phosphorus (P) addition, elevated [CO_2_], nitrogen (N) addition, and their interaction on the foliar traits of invasive and native species as determined by multi-way ANOVA (*F* values are shown in the table). Aarea is photosynthetic rates per unit area; Amass is photosynthetic rate per unit mass; LMA is leaf mass per unit area; PNUE is photosynthetic N-use efficiency and PPUE is photosynthetic P-use efficiency; [N] is N concentration, [P] is P concentration.

| Fixed effect | A<sub>area</sub> | A<sub>mass</sub> | LMA | PNUE | PPUE | [N] | [P] | N:P ratios | Phosphate | Metabolic P | Nucleic acid P | Lipid P | Residual P | Biomass |
| --- | --- | --- | --- | --- | --- | --- | --- | --- | --- | --- | --- | --- | --- | --- |
| S | **39.5** | **134** | **97.5** | **36.4** | 2.8 | **128** | **343** | **16.3** | **212.9** | **57.8** | **190** | **157** | **32.9** | **61.4** |
| P | 0.9 | 1.5 | **9.1** | **14.6** | **32.8** | 2.4 | **55.4** | **55.0** | **13.4** | **14.8** | **10.3** | **57.3** | **3.9** | **14.4** |
| CO<sub>2</sub> | **147** | **41.1** | **5.6** | **36.4** | 2.8 | 1.6 | 3.7 | **11.2** | 0.3 | **8.5** | **15.4** | 1.4 | **6.6** | **27.6** |
| N | **48.0** | **12.1** | 0.1 | 1.4 | **26.0** | **27.4** | 3.4 | **20.4** | 0.6 | 2.7 | 0.02 | 0.4 | 2.5 | **37.0** |
| S × P | 0.6 | 1.1 | **4.4** | 1.4 | **4.5** | **3.8** | **3.7** | **7.6** | 1.6 | **4.5** | 1.3 | **3.9** | **2.9** | 1.7 |
| S × CO<sub>2</sub> | **3.2** | 0.6 | **5.8** | **8.8** | **19.4** | **5.6** | **11.6** | **5.7** | **24.4** | 0.8 | **3.8** | 0.8 | **6.5** | 0.9 |
| CO<sub>2</sub> × P | 1.3 | 1.5 | 0.8 | **6.0** | **9.9** | 0.5 | **5.0** | **6.8** | 2.2 | 2.9 | 2.4 | **5.9** | **2.0** | 2.4 |
| S × P × CO<sub>2</sub> | 1.4 | 1.0 | **4.6** | 2.1 | **2.5** | 0.6 | 0.7 | 1.6 | 1.1 | 1.6 | 1.5 | 1.7 | **4.8** | 0.3 |
| S × P × CO<sub>2</sub> × N | **4.1** | 2.5 | **4.5** | **11.9** | **15.6** | 1.0 | 0.3 | 0.4 | 0.2 | 1.1 | 0.8 | **3.8** | **7.6** | 1.2 |

Significant *F* values (*p*<0.05) are in bold.

**Figure 1.**
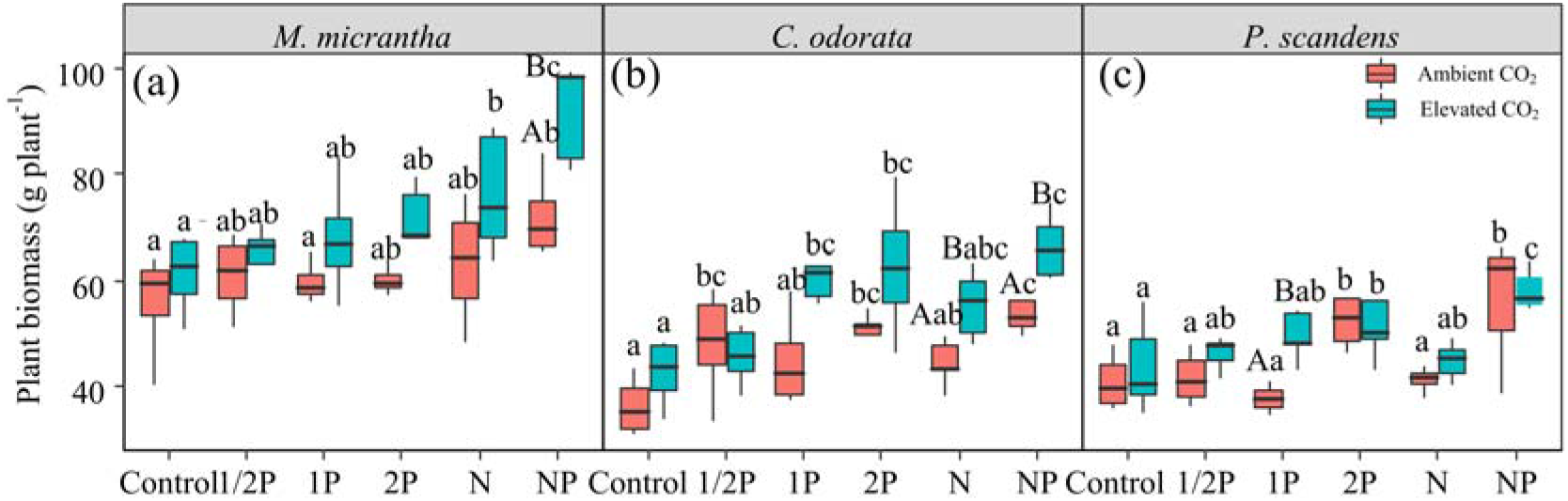
Plant biomass for the invasive species (*Mikania micrantha* and *Chromolaena odorata*) and the native species (*Paederia scandens*) as affected by P addition rate and the combined addition of P and N under ambient or elevated [CO_2_]. The six treatments listed along X axis are as follows: Control (neither P nor N added); 1/2P, 1P, and 2P (0.75, 1.5, and 3 g P m^−2^ yr^−1^, respectively); N (6.25 g N m^−2^ yr^−1^); NP (6.25 g N m^−2^ yr^−1^+1P). For each species and each CO_2_ treatment, means with different lowercase letters are significantly different at *p*<0.05. For each species and within each P or N addition treatment, means with different uppercase letters are significantly different at *p*<0.05. In all cases, the absence of lowercase or uppercase letters indicates the absence of statistical significance.

**Figure 2.**
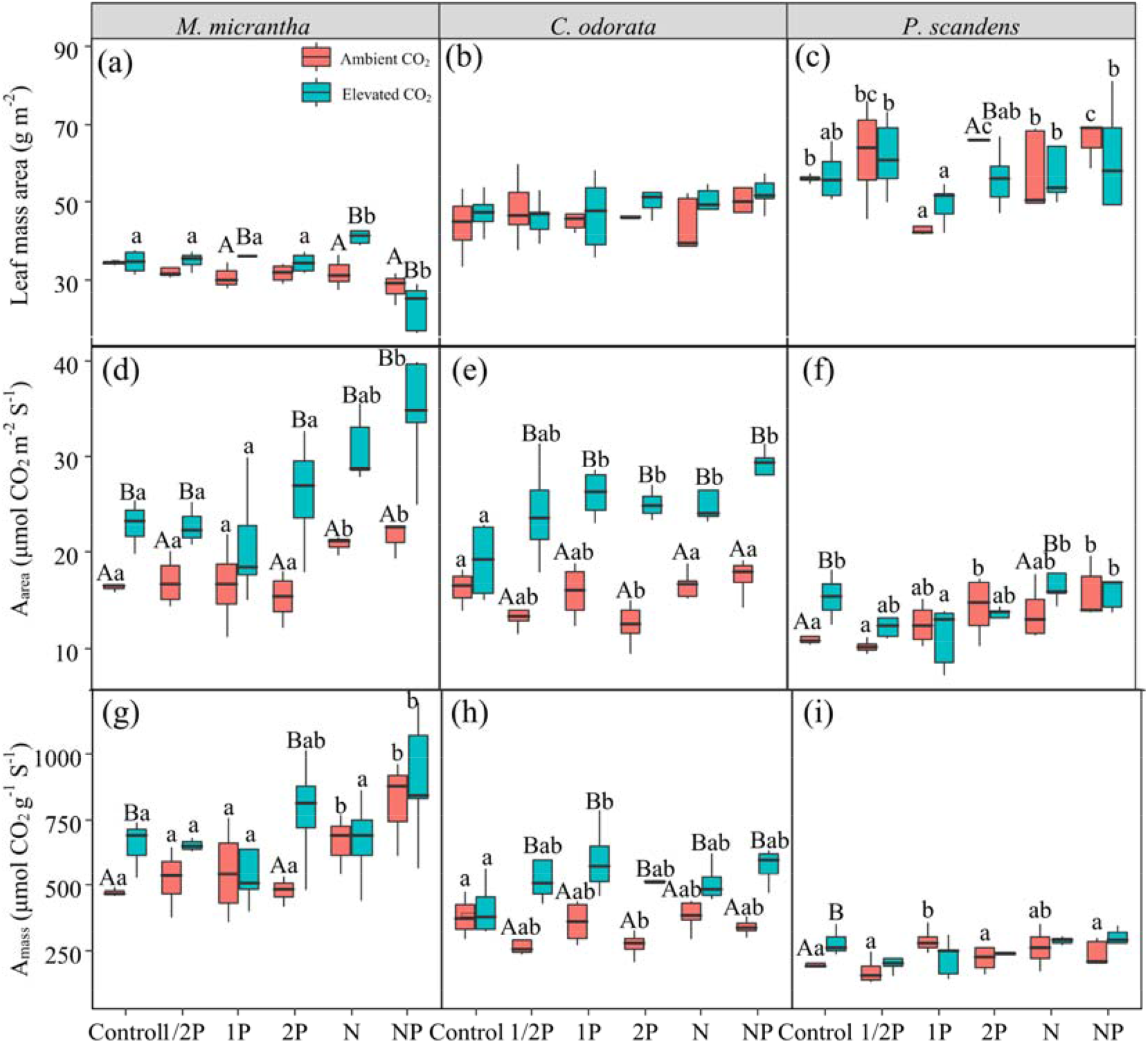
Leaf mass per unit area (LMA), photosynthetic rates per unit area (A_area_), and photosynthetic rates per unit mass (A_mass_) for the invasive species (*M. micrantha* and *C. odorata*) and the native species (*P. scandens*) as affected by P addition rate and the combined addition of P and N under ambient or elevated [CO_2_]. Treatments and statistical comparisons are described in Figure 1.

### PPUE and PNUE

The interaction of species, elevated [CO_2_], N addition, and P addition significantly affected PNUE and PPUE (Table 1). On average, elevated [CO_2_] increased PNUE and PPUE by 62% and 51%, respectively, in *C. odorata*, and by 79% and 41%, respectively, in *P. scandens* (Fig. 3b, c, e, f). Elevated [CO_2_] significantly increased PPUE in *M. micrantha* (by 73%), but only when combined with 2P addition (Fig. 3d). N addition did not significantly affect PNUE in the invasive species, but increased PPUE in *M. micrantha* (Fig. 3a, b, d, e). Under elevated [CO_2_], 1/2P, 1P and 2P addition significantly increased PNUE by 24%, 55% and 57%, respectively, in *C. odorata*, and 2P addition significantly increased PNUE by 34% in *P. scandens* (Fig. 3a, b, c). However, PPUE significantly decreased in all species as the quantity of P added was increased further. 2P addition significantly decreased PPUE by 25% in *C. odorata* and by 32% in *P. scandens* under elevated [CO_2_] (Fig. 3d, e, f).

**Figure 3.**
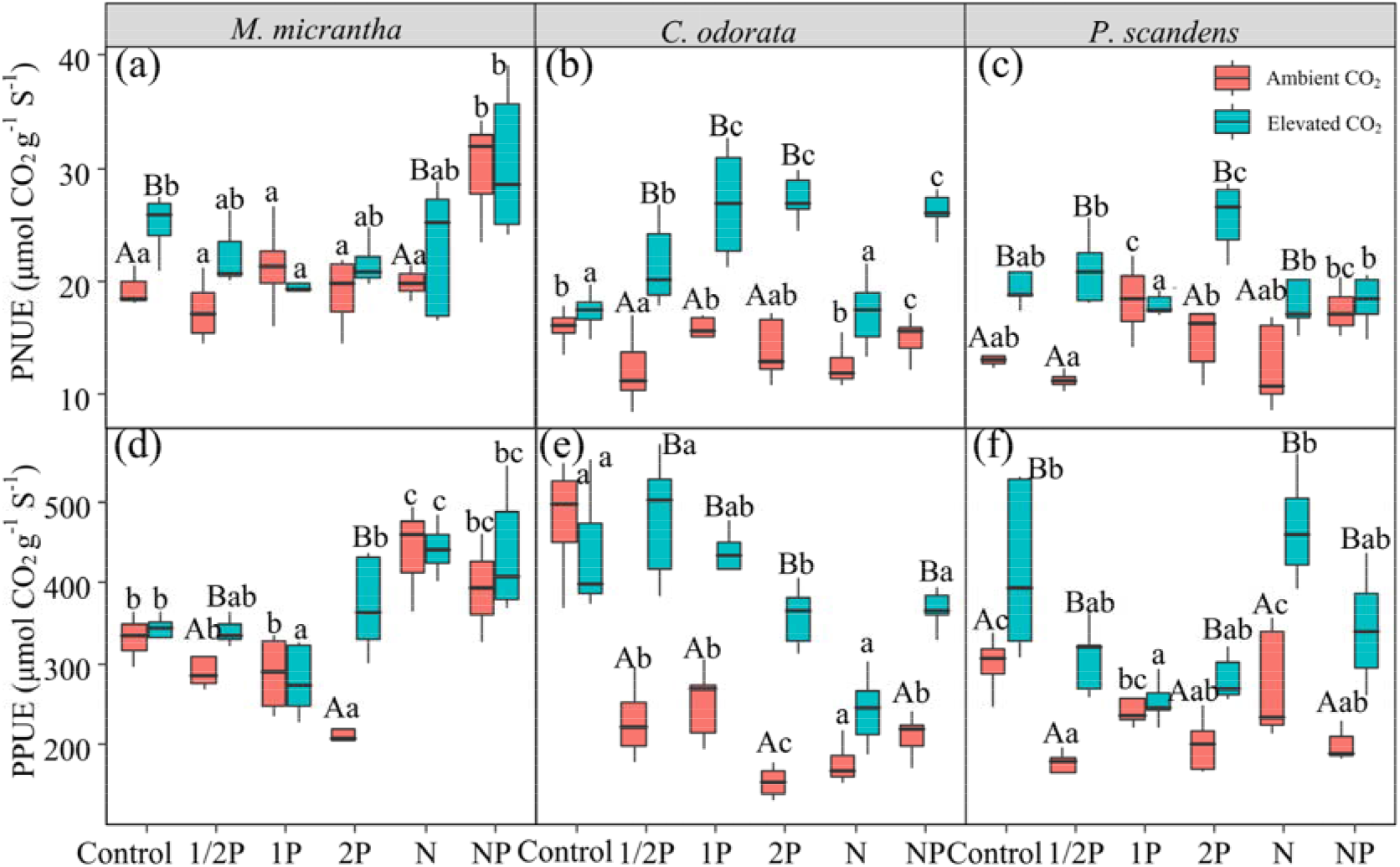
Photosynthetic nutrient-use efficiency for nitrogen (PNUE) and phosphorus (PPUE) for invasive species *M. micrantha* and *C. odorata* and the native species *P. scandens* as affected by P addition rate and the combined addition of P and N under ambient or elevated [CO_2_]. Treatments and statistical comparisons are described in Figure 1.

### Foliar P and N

Species and its interaction with elevated [CO_2_], N addition, and P addition significantly affected foliar P and N concentrations (Table 1). N addition significantly increased the foliar N concentration, but slightly decreased the foliar P concentration (Fig. 4). Under elevated [CO_2_], 2P addition significantly decreased N concentrations by 15% in *C. odorata* and by 13% in *P. scandens* (Fig. 4a, b, c). Foliar P concentrations significantly increased with increases in the amount of P added in *M. micrantha* and *C. odorata,* and the increases were greater for the invasive species than for *P. scandens*; this effect of added P was slightly reduced by elevated [CO_2_]. 2P addition significantly increased foliar P concentrations by 50 and 96% in *M. micrantha* and *C. odorata*, respectively, and by 58.3% in *P. scandens* (Fig. 4d, e, f).

**Figure 4.**
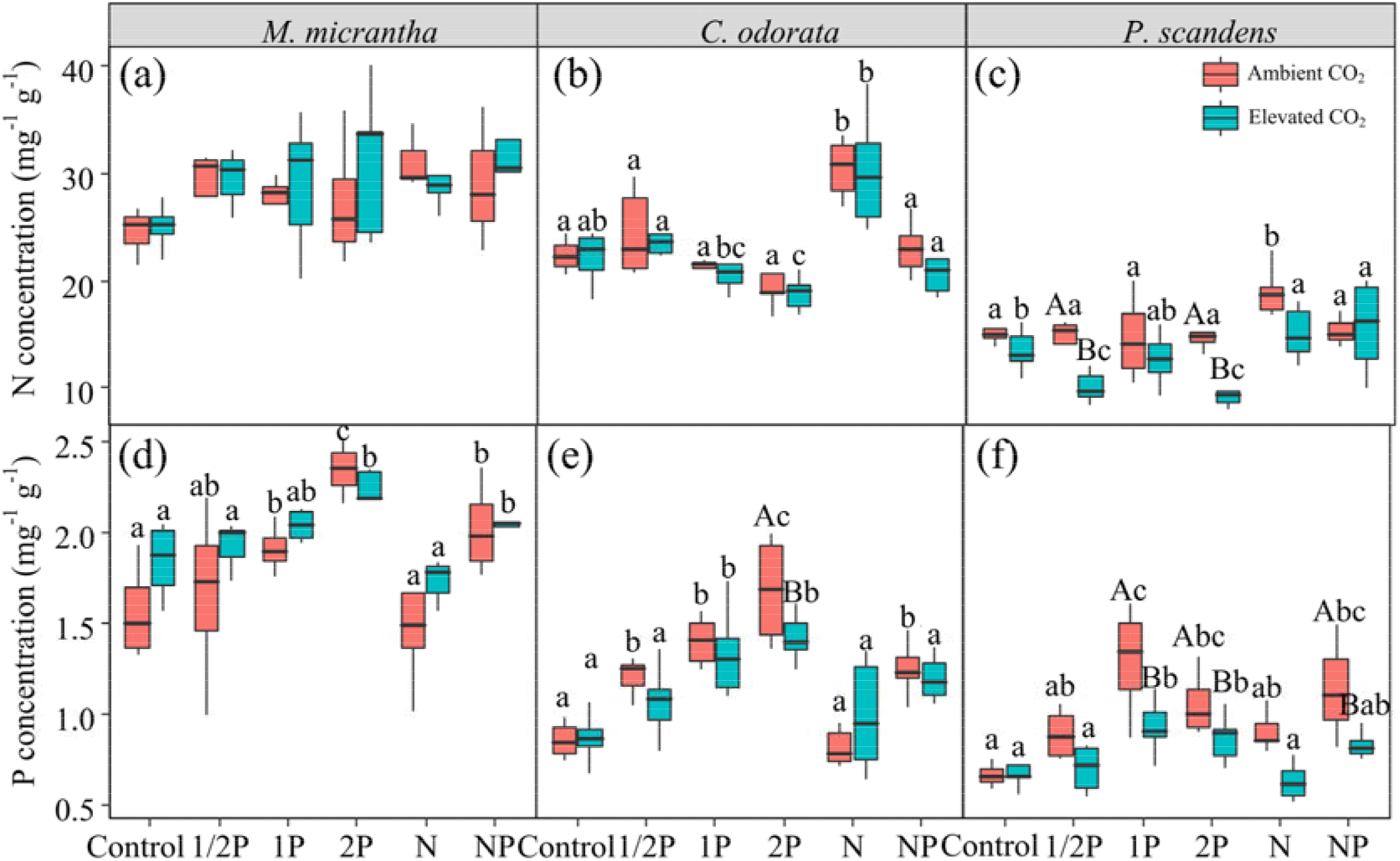
Foliar nitrogen (N) and phosphorus (P) concentrations of the invasive species *M. micrantha* and *C. odorata* and the native species *P. scandens* as affected by P addition rate and the combined addition of P and N under ambient or elevated [CO_2_]. Treatments and statistical comparisons are described in Figure 1.

### Foliar P fractions and N:P ratios

Overall, both species and P addition significantly affected foliar P fractions, i.e. Pi, metabolic P, nucleic P, lipid P and residual P (Table 1). N addition slightly decreased the concentrations of all foliar P fractions in all species (Fig. 5). In *M. micrantha* and *C. odorata*, Pi (Fig. 5a, b), metabolic P (Fig. 5d, e), nucleic acid P (Fig. 5g, h) and lipid P (Fig. 5j, k) were significantly increased by an average of 53, 754, 38, and 82%, respectively, by 2P addition, but residue P was not significantly increased by P addition (Fig. 5m, n). In *P. scandens*, foliar P fractions significantly increased with the amount of P added from 1/2P to 1P, but tended to decrease with 2P addition (Fig. 5f, i, l, o). In response to P addition, metabolic P increased the most, followed by nucleic acid P, Pi, and lipid P in all species, while residual P did not respond in a consistent pattern to P addition. Elevated [CO_2_] slightly weakened the enhancing effect of P addition on foliar P fractions in all species (Fig. 5). The concentration of foliar Pi, metabolic P, nucleic P, lipid P and residual P are all significantly and positively correlated with plant biomass, suggested that plant biomass increased with the increase of the concentration of foliar P fractions (Fig. 6).

**Figure 5.**
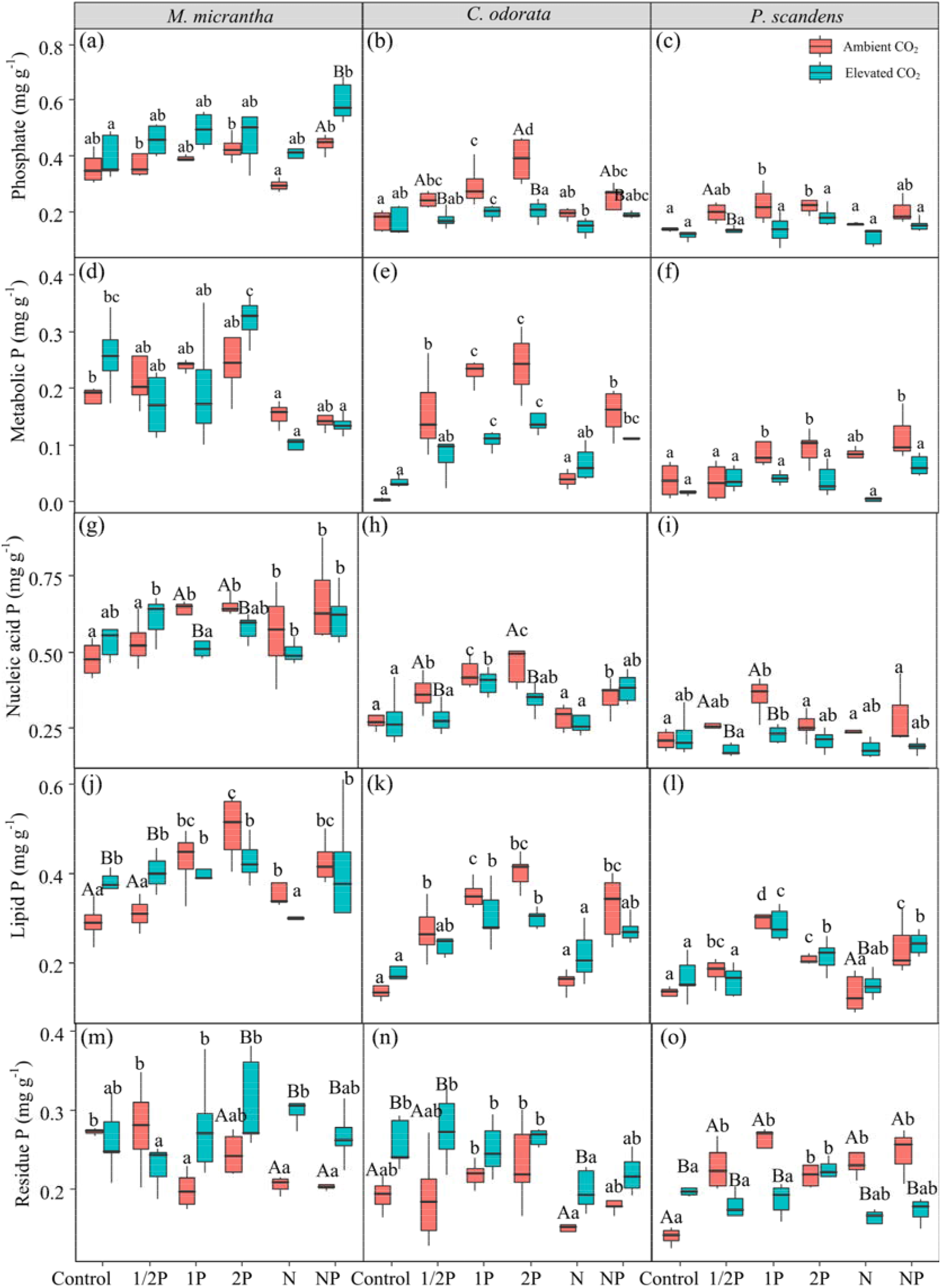
Concentrations of foliar inorganic P (phosphate, Pi) and organic P (metabolic P, nucleic acid P, structural P, and residual P) of the invasive species *M*. *micrantha* and *C. odorata* and the native species *P. scandens* as affected by P addition rate and the combined addition of P and N under ambient or elevated [CO_2_]. Treatments and statistical comparisons are described in Figure 1.

**Figure 6.**
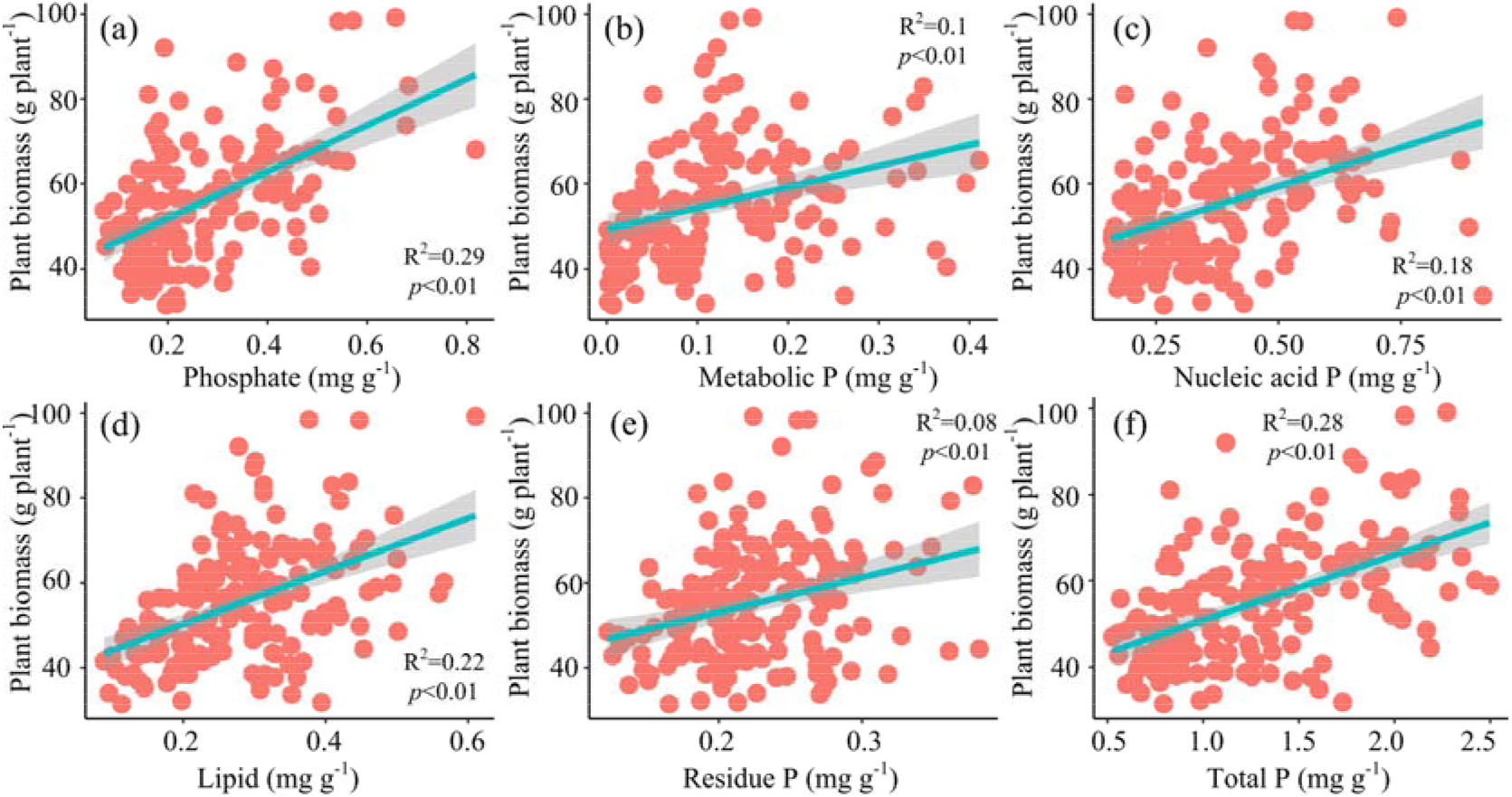
Correlation between foliar phosphorus (P) fractions concentration and plant biomass. The data from three target species and six treatments. *R*^2^ values for linear trend lines are shown on each plot.

As the amount of P added was increased, the foliar N: P ratios, foliar N: phosphate ratios, foliar N: metabolic P ratios and foliar N: lipid P ratios decreased more in *M. micrantha* and *C. odorata* than in *P. scandens* under ambient [CO_2_] (Table 2). Under ambient [CO_2_], however, the foliar N: residue P ratio did not significantly differ among 1/2P, 1P, and 2P treatments in any species, except between 1/2P and 1P in *C. odorata* (Table 2). Elevated [CO_2_] significantly decreased the foliar N: phosphate ratio, foliar N: lipid P ratio, and foliar N: residue P ratio in *C. odorata* (Table 2). In the invasive species, the concentrations of foliar P fractions (phosphate, metabolic P, nucleic P, lipid P and residual P) were positively correlated with A_mass_, N concentration, and P concentration, but negatively with LMA, PPUE, and N: P ratios (Table 3).

**Table 2.**
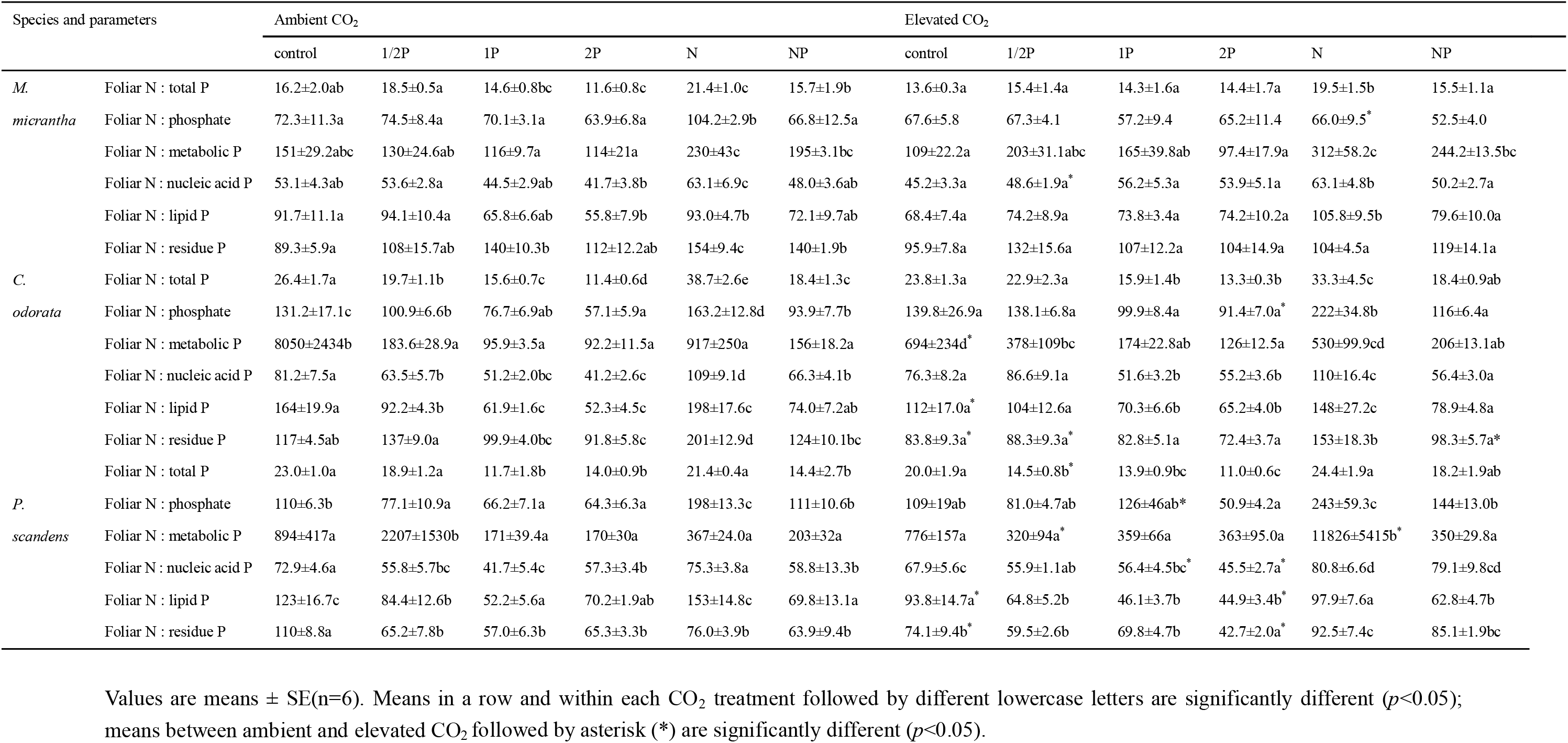
Ratios of total foliar nitrogen (N) to phosphorus (P) in P-containing leaf fractions.

| Species and parameters |  | Ambient CO<sub>2</sub> |  |  |  |  |  | Elevated CO<sub>2</sub> |  |  |  |  |  |
| --- | --- | --- | --- | --- | --- | --- | --- | --- | --- | --- | --- | --- | --- |
|  |  | control | 1/2P | 1P | 2P | N | NP | control | 1/2P | 1P | 2P | N | NP |
| *M.* | Foliar N : total P | 16.2±2.0ab | 18.5±0.5a | 14.6±0.8bc | 11.6±0.8c | 21.4±1.0c | 15.7±1.9b | 13.6±0.3a | 15.4±1.4a | 14.3±1.6a | 14.4±1.7a | 19.5±1.5b | 15.5±1.1a |
| *micrantha* | Foliar N : phosphate | 72.3±11.3a | 74.5±8.4a | 70.1±3.1a | 63.9±6.8a | 104.2±2.9b | 66.8±12.5a | 67.6±5.8 | 67.3±4.1 | 57.2±9.4 | 65.2±11.4 | 66.0±9.5* | 52.5±4.0 |
|  | Foliar N : metabolic P | 151±29.2abc | 130±24.6ab | 116±9.7a | 114±21a | 230±43c | 195±3.1bc | 109±22.2a | 203±31.1abc | 165±39.8ab | 97.4±17.9a | 312±58.2c | 244.2±13.5bc |
|  | Foliar N : nucleic acid P | 53.1±4.3ab | 53.6±2.8a | 44.5±2.9ab | 41.7±3.8b | 63.1±6.9c | 48.0±3.6ab | 45.2±3.3a | 48.6±1.9a* | 56.2±5.3a | 53.9±5.1a | 63.1±4.8b | 50.2±2.7a |
|  | Foliar N : lipid P | 91.7±11.1a | 94.1±10.4a | 65.8±6.6ab | 55.8±7.9b | 93.0±4.7b | 72.1±9.7ab | 68.4±7.4a | 74.2±8.9a | 73.8±3.4a | 74.2±10.2a | 105.8±9.5b | 79.6±10.0a |
|  | Foliar N : residue P | 89.3±5.9a | 108±15.7ab | 140±10.3b | 112±12.2ab | 154±9.4c | 140±1.9b | 95.9±7.8a | 132±15.6a | 107±12.2a | 104±14.9a | 104±4.5a | 119±14.1a |
| *C.* | Foliar N : total P | 26.4±1.7a | 19.7±1.1b | 15.6±0.7c | 11.4±0.6d | 38.7±2.6e | 18.4±1.3c | 23.8±1.3a | 22.9±2.3a | 15.9±1.4b | 13.3±0.3b | 33.3±4.5c | 18.4±0.9ab |
| *odorata* | Foliar N : phosphate | 131.2±17.1c | 100.9±6.6b | 76.7±6.9ab | 57.1±5.9a | 163.2±12.8d | 93.9±7.7b | 139.8±26.9a | 138.1±6.8a | 99.9±8.4a | 91.4±7.0a* | 222±34.8b | 116±6.4a |
|  | Foliar N : metabolic P | 8050±2434b | 183.6±28.9a | 95.9±3.5a | 92.2±11.5a | 917±250a | 156±18.2a | 694±234d* | 378±109bc | 174±22.8ab | 126±12.5a | 530±99.9cd | 206±13.1ab |
|  | Foliar N : nucleic acid P | 81.2±7.5a | 63.5±5.7b | 51.2±2.0bc | 41.2±2.6c | 109±9.1d | 66.3±4.1b | 76.3±8.2a | 86.6±9.1a | 51.6±3.2b | 55.2±3.6b | 110±16.4c | 56.4±3.0a |
|  | Foliar N : lipid P | 164±19.9a | 92.2±4.3b | 61.9±1.6c | 52.3±4.5c | 198±17.6c | 74.0±7.2ab | 112±17.0a* | 104±12.6a | 70.3±6.6b | 65.2±4.0b | 148±27.2c | 78.9±4.8a |
|  | Foliar N : residue P | 117±4.5ab | 137±9.0a | 99.9±4.0bc | 91.8±5.8c | 201±12.9d | 124±10.1bc | 83.8±9.3a* | 88.3±9.3a* | 82.8±5.1a | 72.4±3.7a | 153±18.3b | 98.3±5.7a* |
|  | Foliar N : total P | 23.0±1.0a | 18.9±1.2a | 11.7±1.8b | 14.0±0.9b | 21.4±0.4a | 14.4±2.7b | 20.0±1.9a | 14.5±0.8b* | 13.9±0.9bc | 11.0±0.6c | 24.4±1.9a | 18.2±1.9ab |
| *P.* | Foliar N : phosphate | 110±6.3b | 77.1±10.9a | 66.2±7.1a | 64.3±6.3a | 198±13.3c | 111±10.6b | 109±19ab | 81.0±4.7ab | 126±46ab* | 50.9±4.2a | 243±59.3c | 144±13.0b |
| *scandens* | Foliar N : metabolic P | 894±417a | 2207±1530b | 171±39.4a | 170±30a | 367±24.0a | 203±32a | 776±157a | 320±94a* | 359±66a | 363±95.0a | 11826±5415b* | 350±29.8a |
|  | Foliar N : nucleic acid P | 72.9±4.6a | 55.8±5.7bc | 41.7±5.4c | 57.3±3.4b | 75.3±3.8a | 58.8±13.3b | 67.9±5.6c | 55.9±1.1ab | 56.4±4.5bc* | 45.5±2.7a* | 80.8±6.6d | 79.1±9.8cd |
|  | Foliar N : lipid P | 123±16.7c | 84.4±12.6b | 52.2±5.6a | 70.2±1.9ab | 153±14.8c | 69.8±13.1a | 93.8±14.7a* | 64.8±5.2b | 46.1±3.7b | 44.9±3.4b* | 97.9±7.6a | 62.8±4.7b |
|  | Foliar N : residue P | 110±8.8a | 65.2±7.8b | 57.0±6.3b | 65.3±3.3b | 76.0±3.9b | 63.9±9.4b | 74.1±9.4b* | 59.5±2.6b | 69.8±4.7b | 42.7±2.0a* | 92.5±7.4c | 85.1±1.9bc |

Values are means ± SE(n=6). Means in a row and within each CO<sub>2</sub> treatment followed by different lowercase letters are significantly different ( $p<0.05$ ); means between ambient and elevated CO<sub>2</sub> followed by asterisk (\*) are significantly different ( $p<0.05$ ).

**Table 3.** Correlations between foliar phosphorus (P) fractions and foliar traits in the two invasive species. A_area_ is photosynthetic rates per unit area; A_mass_ is photosynthetic rates per unit mass; LMA is leaf mass per unit area; PNUE is photosynthetic N-use efficiency; PPUE is photosynthetic P-use efficiency; [N] is N concentration, [P] is P concentration. Values are correlation coefficients. * and ** indicates significance at *p*<0.05 and *p*<0.01, respectively.

| P fraction | $A_{\text{area}}$ | $A_{\text{mass}}$ | LMA | PNUE | PPUE | [N] | [P] | N:P ratios |
| --- | --- | --- | --- | --- | --- | --- | --- | --- |
| Phosphate | 0.33** | 0.58** | -0.55** | 0.24** | 0.01 | 0.47** | 0.80** | -0.36** |
| Metabolic P | 0.12 | 0.32* | -0.42** | 0.03 | -0.13 | 0.39** | 0.70** | -0.32** |
| Nucleic acid P | 0.35** | 0.61** | -0.61** | 0.26** | -0.05 | 0.66** | 0.88** | -0.24** |
| Lipid P | 0.29** | 0.53** | -0.59** | 0.30** | -0.20* | 0.49** | 0.89** | -0.43** |
| Residual P | 0.36* | 0.39** | -0.20* | 0.28** | -0.04 | 0.29** | 0.54** | -0.30** |

## Discussion

Phosphorus addition significantly increased the plant biomass of *M. micrantha* and *C. odorata*, which was consistent with previous studies that P limitation of plant growth occurred in subtropical forest ecosystems (Hidaka and Kitayama, 2013; Hou et al., 2020). Recent studies, however, the invasive species *M. micrantha* and *C. odorata* in a soil with low P availability under ambient conditions of [CO_2_] and N deposition still maintain high plant biomass, and rapid invasion in the forest ecosystem (Tang et al., 2007; Song et al., 2009; Zhang et al., 2016). In the present study, we found that changes in allocation of foliar P fractions after P addition which may help explain how invasive species can maintain their photosynthetic capacity and therefore possibly their invasiveness in soils with low P availability under conditions of elevated atmospheric [CO_2_] and N deposition.

### P addition affected foliar traits and photosynthetic capacity under the combination of elevated [CO_2_] and N addition

Although we observed no significant increase of foliar N in response to P addition in any of the studied species, we found a marked increase in response to P addition in foliar P concentration and plant biomass in the two invasive species, and this increase was greater than that in the native species. These results indicated that P addition alleviated the P limitation of plant growth (Buenemann et al., 2011; Novriyanti et al., 2012). However, N and P addition did not significantly affect photosynthesis under ambient [CO_2_], including that photosynthesis were not limited by N or P in ambient [CO_2_] (Dissanayaka et al., 2018). Elevated [CO_2_] did not significantly affect LMA or concentrations of foliar N or P, The most likely explanation for this is that plants that experienced increased A_area_ and A_mass_ under elevated [CO_2_] could not use the newly fixed carbohydrates for new growth (Presoctt et al., 2020). In our study, A_area_ and A_mass_ increased with an increase in P supply under elevated [CO_2_] and N addition in the invasive species, consistent with previous studies that elevated [CO_2_] and N deposition generally increased photosynthetic capacity (A_area_ and A_mass_) even in low-P soils (Campbell and Sage, 2006), and consequently increased plant growth (Zhang et al., 2016). P addition can alleviate P limitation for plant growth and can increase photosynthesis under elevated [CO_2_], consistent with our first two hypotheses. With increasing soil P availability under elevated [CO_2_], the invasive plants showed a remarkable increase in foliar P concentrations, and provided enough metabolic P for intermediates of nucleotide and carbon metabolism for photosynthetic rates (Hidaka and Kitayama, 2013; Mo et al., 2019). We also found that the interaction of N and P addition under elevated [CO_2_] increased the photosynthesis to a greater degree in invasive species than in the native species.

Plants in P-deficient soils also change foliar PNUE and PPUE to maintain their growth (Zhang et al., 2016; Mo et al., 2019). In our study, the interaction of P addition and elevated [CO_2_] significantly increased PNUE, and N addition did not significantly increased plant biomass of all species, which both indicated that the soil even without N addition contained sufficient N to support increased growth. In our study, PNUE was higher in invasive than in native species despite insignificant differences in foliar N concentration in treatments that differed in quantities of P added under elevated [CO_2_], probably due to the higher photosynthetic capacity (A_area_ and A_mass_) of invasive species than native species. We also found that PPUE in invasive species greatly decreased with increasing rates of P addition, but increased with N addition and elevated [CO_2_], indicating that plants experiencing low levels of P use P more efficiently than plants experiencing adequate levels of P, and that soil P availability affected PPUE more in invasive species than in native species. The results were in agreement with a previous study, which found that PPUE decreased in response to increasing soil P availability, because the accumulation of P in plants was not adequately be used (Hidaka and Kitayama, 2011).

### P addition affected foliar P fractions under the combination of elevated [CO_2_] and N addition

In our study, the foliar N: total P and N: P fractions ratios were >20, and plant biomass increased with the increasing amount of P application in the invasive species, indicating that P limitation occurred in untreated plants (Güsewell, 2004; Tang et al., 2007; Mo et al., 2019). P addition increased plant biomass by increasing foliar P fractions (Pi, metabolic P, lipid P and nucleic acid P), and decreased foliar N:P ratios, which might promote the photosynthesis and growth of invasive species by increasing the amount of foliar P available for synthesis of rRNA and membrane phospholipids (Reef et al., 2010).

Changes in the foliar N:P ratio has previously been associated with physiological growth strategies in both invasive and native species (Hidaka and Kitayama, 2011). Alleviation of P addition was greater in the two invasive species than in the native species and involved increases in foliar P concentrations in the invasive species. This effect, however, was weakened by elevated [CO_2_] and N addition, which was consistent with a previous finding that elevated [CO_2_] and N addition increased the foliar N: P ratio and exacerbated plant P limitation (Zhang et al., 2016). Therefore, P addition significantly decreased foliar N:P ratios, and increased plant biomass.

The shifts in the foliar P fractions with increasing amounts of P addition under conditions of elevated [CO_2_] and N addition also provide clues to the underlying adaptive mechanisms that explain the success of invasive species (Hidaka and Kitayama, 2011). Invasive plants may regulate the balance between the levels of phosphorylated intermediates and inorganic P in order to maintain photosynthetic rates when soil P availability is deficient (Wang et al., 2019). In our study, although P addition increased the concentrations of P fractions concentrations of invasive species, P addition did not affect the LMA. These results indicate that P addition greatly changed foliar P allocation, rather than LMA so as to maintain stable photosynthetic rates when plant growth is limited by phosphorus. Plants store a large amount of Pi in their vacuoles, which could provide sufficient triose phosphates for chloroplasts and photophosphorylation for plant photosynthesis (Mo et al., 2019). In the current study, P addition significantly increased foliar Pi (the largest proportion of foliar P) for *M. micrantha* and *C. odorata*; among foliar P fractions, the increase was not highest for leaf Pi, because Pi in the leaf is generally diverted to other P fractions for photosynthesis to meet the demand of plant growth (Ostertag, 2010). The concentration of the metabolic P fraction in all species was low but increased more than the other P fractions in response to P addition. The largest increase in metabolic P indicated that plant metabolic intermediates (e.g., phytate) increased in response to P addition, which consequently increased carbon metabolism and nucleotides (Ostertag, 2010; Veneklaas et al., 2012).

Consistent with our third hypothesis, the increases in Pi and metabolic P fractions with the increasing amounts of P addition were greater in the invasive species than in native species, but elevated [CO_2_] and N addition slightly weakened this effect. A possible explanation is that the Pi and metabolic P fractions are transformed into membrane phospholipids under P addition, and that this transformation is weakened by elevated [CO_2_] and N addition (Lambers et al., 2015). Previous studies found that the status of Pi in the cytosol strongly affected photosynthetic rates (Schachman et al., 1998), and that the interaction of elevated [CO_2_] and N addition supported stable photosynthetic rates by maintaining Pi and metabolic P in invasive species in P-poor soils (Lambers et al., 2015). Other studies have reported that, to maintain photosynthesis when soil P availability is low, plants can use P from lipids or nucleic acids to maintain foliar P in the form of Pi and metabolic P in the cytosol (Ostertag, 2010; Mo et al., 2019; Prodhan et al., 2019). In contrast, we observed that P addition increased the concentrations of all foliar P fraction, and that the increases were greatest for metabolic P and Pi in plant cell. These results further indicate that invasive plants can alter the balance between foliar Pi or metabolic P and other fractions (lipid P and nucleic P) in order to maintain a high photosynthetic capacity in soils with low P availability.

### Relationships between foliar traits and P fractions

In our study, the concentrations of foliar P fractions (Pi, metabolic P, lipid P, and nucleic acid P) were negatively correlated with LMA under conditions of different levels of P, N, and [CO_2_]. When growing in soil with low P availability in subtropical forest, P-limited invasive plants develop thick and tough leaves with a high LMA in order to prolong leaf life span (Ellsworth and Reich, 1996; Hidaka and Kitayama, 2011). Increased LMA may lead to a decrease in photosynthetic capacity (A_area_ and A_mass_) because of the increased resistance to CO_2_ diffusion. In the current study, the decrease in the foliar P concentration for invasive plants growing in soil without P addition was associated with an increase in LMA and seemed to indicate a reduced demand for P. PPUE was negatively correlated with structural P for the invasive species growing in soil with low P availability, which indicated that the invasive species were able to slightly increase LMA without sacrificing PPUE by decreasing concentrations of foliar P fractions (Pi, metabolic P, lipid P, and nucleic acid P), as suggested by Mo et al., (2019).

The results in this study were used to develop a conceptual framework for the mechanism of P maintenance in low P soil availability under elevated CO_2_ and N addition (Fig. 7). Elevated CO_2_ and N addition exacerbated the P demand of invasive species, which was indicated by increased ratios of foliar N:P and foliar N: fractions P, and decreased foliar P concentration after elevated CO_2_ and N addition. Therefore, the transformation of non-metabolic P (lipid P and nucleic acid P) to metabolic P and phosphate was enhanced. This pathway was essential to meet the increased P requirement for the growth of invasive species in low soil P availability under elevated CO_2_ and N addition in the subtropical forest ecosystem.

**Figure 7.**
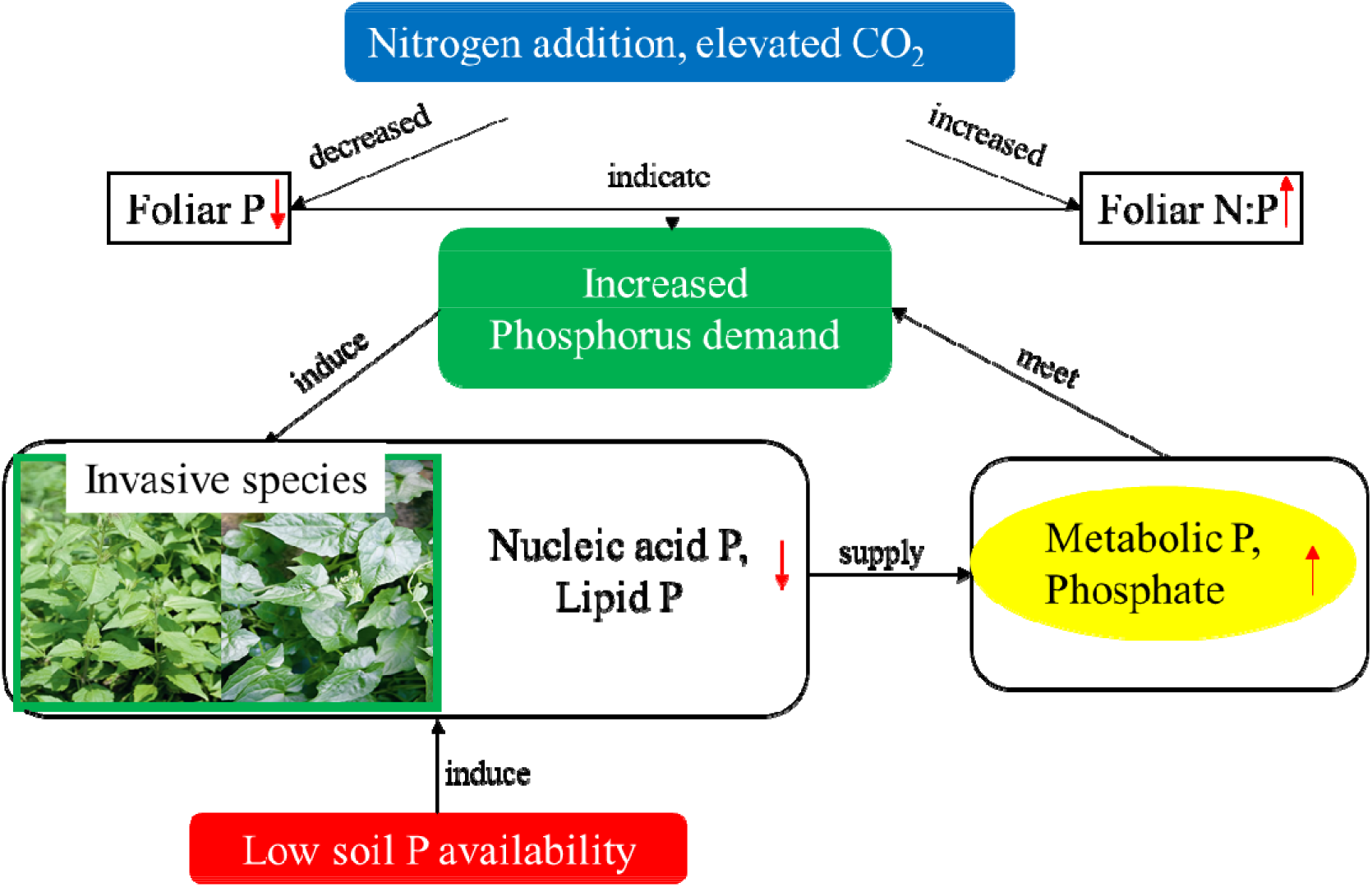
A conceptual framework of a noval pathway to sustain P demand of invasive species under elevated CO_2_ and N addition in low soil P availability.

## Conclusions

In the current study, elevated [CO_2_] more than N addition allowed invasive plants to adjust their foliar traits and acclimate to low soil P availability; the acclimation was substantially greater in the two invasive species than in the native species. Plant biomass significantly increased under P addition, and the foliar N: P ratio >20 of the invasive species indicated P limitation of its growth. Rather than decreasing their LMA, the invasive species acclimated to low soil P availability under elevated [CO_2_] and N addition by greatly reducing their allocation of P to non-metabolic foliar P fractions (nucleic acid P and lipid P); conversely metabolic P and Pi were not reduced, and this may have allowed maintenance of a high photosynthetic capacity. These adaptive responses help explain the success of invasive plants under conditions of rising atmospheric [CO_2_] and N deposition in soil with low P availability. This knowledge may prevent them from rapid proliferation with global climate changes.

## Acknowledgments

This research was funded by the National Natural Science Foundation of China (31570401; 31100411), the Key Special Project for Introduced Talents Team of Southern Marine Science and Engineering Guangdong Laboratory (Guangzhou) (GML2019ZD0408), Postdoctoral Science Foundation of China (2020M682950), and the NSFC-Guangdong Joint Fund, China (U1701246).

## Author’s contributions

L.Z. and D.W. conceived and designed the research; L.Z., G.Z., N.L., X.Z. and M.X. performed the research; L.Z. and X.L. analyzed and interpreted the data; L.Z. and X.L. wrote the paper, H.L. revised the manuscript.

---

**Figure S1.**
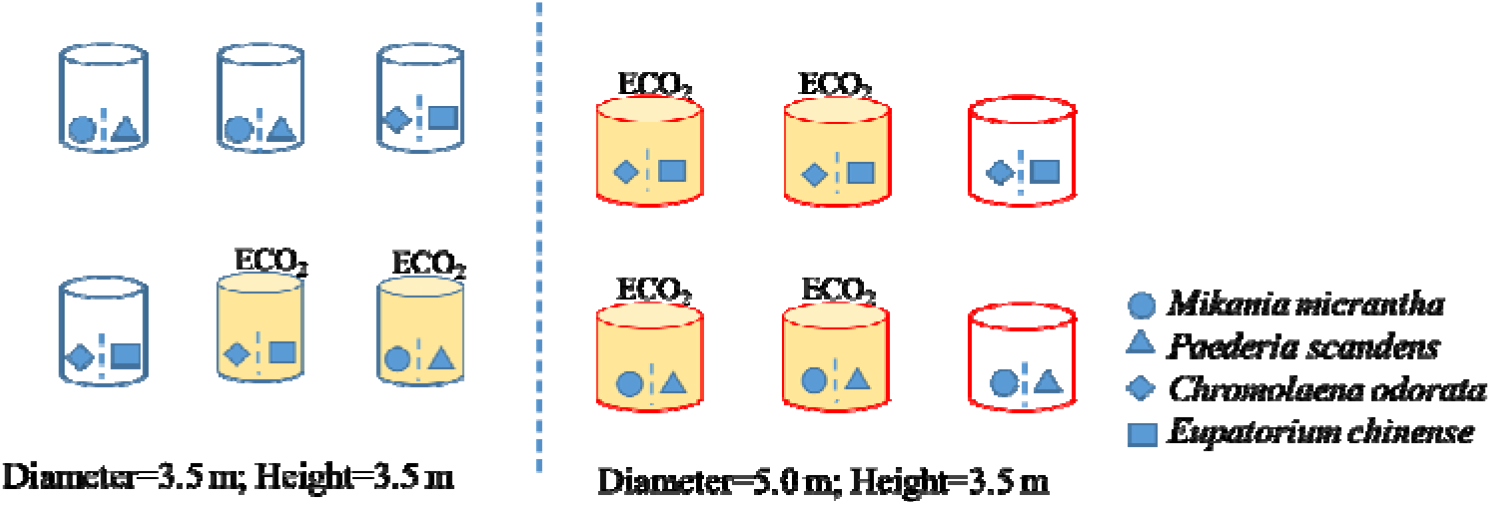
Diagram of species planted in open-top chambers. Chambers labeled ECO_2_ were treated with elevated [CO_2_]. The other chambers were treated with ambient [CO_2_]. Introduce: To clearly discriminate the response of invasive species to the joint effects, two indigenous co-occurring species (*Paederia scandens* and *Eupatorium chinense*) with similar morphology to invasive species were together collected. *Mikania micrantha*, *Chromolaena odorata* and *P. scandens* were collected in South China Botanical Garden, *E. chinense* was collected in Zhejiang Province. Most of the treated *E. chinense* dead in the cultivating process, as they could not adapt to the climate conditions in Guangzhou, China. So the foliar parameters of *E. chinense* couldn’t be measured, and no data showed in the paper.

